# Improving spatial normalization of brain diffusion MRI to measure longitudinal changes of tissue microstructure in the cortex and white matter

**DOI:** 10.1101/590521

**Authors:** Florencia Jacobacci, Jorge Jovicich, Gonzalo Lerner, Edson Amaro, Jorge L. Armony, Julien Doyon, Valeria Della-Maggiore

## Abstract

**Background:** Fractional anisotropy (*FA*) and mean diffusivity (*MD*) are frequently used to evaluate longitudinal changes in white matter microstructure. Recently, there has been a growing interest in identifying experience-dependent plasticity in gray matter using *MD*. Improving registration has thus become a major goal to enhance the detection of subtle longitudinal changes in cortical microstructure.

**Purpose:** To optimize normalization to improve registration in gray matter and reduce variability associated with multi-session registrations.

**Study Type:** Prospective longitudinal study

**Subjects:** Twenty-one healthy subjects (18-31 years old) underwent 9 magnetic resonance imaging (MRI) scanning sessions each.

**Field Strength/Sequence:** 3.0T, diffusion-weighted multiband-accelerated sequence, MP2RAGE sequence.

**Assessment:** Diffusion-weighted images were registered to standard space using different pipelines that varied in the features used for normalization, namely the non-linear registration algorithm (FSL vs ANTs), the registration target (*FA*-based vs T1-based templates), and the use of intermediate individual (FA-based or T1-based) targets. We compared the across-session test-retest reproducibility error from these normalization approaches for *FA* and *MD* in white and gray matters.

**Statistical Tests:** Reproducibility errors were compared using a repeated-measures analysis of variance with pipeline as within-subject factor.

**Results:** The registration of FA data to the FMRIB58 FA atlas using ANTs yielded lower reproducibility errors in white matter (p<0.0001) with respect to FSL. Moreover, using the MNI152 T1 template as the target of registration resulted in lower reproducibility errors for *MD* (p<0.0001), whereas the FMRIB58 FA template performed better for *FA* (p<0.0001). Finally, the use of an intermediate individual template improved reproducibility when registration of the FA images to the MNI152-T1 was carried out within modality (FA-FA) (p<0.05), but not via a T1-based individual template.

**Data Conclusion:** A normalization approach using ANTs to register FA images to the MNI152 T1 template via an individual FA template minimized test-retest reproducibility errors both for gray and white matter.

## INTRODUCTION

Due to the growing interest in identifying neuroplasticity induced by learning, scalar DTI measures such as fractional anisotropy (*FA*) and mean diffusivity (*MD*) are being increasingly used in humans to evaluate longitudinal changes in brain tissue microstructure. In healthy individuals, *FA* has been used as an indirect measure of myelination to detect plasticity in white matter (WM) tracts (1, 2). In contrast, *MD* is also sensitive to microstructural changes in gray matter (GM), thereby offering a great potential as a marker of cortical plasticity (3, 4). Indeed, previous studies have shown that learning induces a subtle (~3%) but consistent reduction in MD that reflects astrocyte hypertrophy, possibly triggered by long-term potentiation (4).

When attempting to identify such subtle signal variations in gray matter, the specific image pre-processing approach becomes critical as it may help true signal detection, by reducing noise, or hinder it, by introducing artifacts. Yet, the tools currently available, including the Tract-Based Spatial Statistics (TBSS) algorithm (5, 6), are typically optimized to detect differences in white matter microstructure but not in gray matter. Another important challenge encountered when using DTI to detect longitudinal changes relates to the possible occurrence of registration errors resulting from the alignment of multiple images to a standard stereotaxic space (5, 7). To date, however, there is no consensus regarding the optimal normalization pipeline to process longitudinal DTI data (8).

The overall goal of this study was to generate a normalization pipeline to optimize registration of longitudinal DTI data in gray matter. To this aim, we systematically compared the effect of a combination of features in the normalization process of diffusion images to minimize across-session test-retest reproducibility errors (RE) in a group of healthy volunteers scanned over repeated sessions. More specifically, we tested the impact of three features: 1) the registration algorithm used for normalization (FSL vs ANTs), 2) the target template used as the reference in the normalization process (*FA*-based vs MNI152 templates), and 3) the use of intermediate targets, i.e., different intermediate individual templates (*FA*-based or T1-based) to register to the stereotaxic space. Our working hypothesis was that minimizing the test-retest reproducibility error of DTI measures would help improve the detection of subtle longitudinal changes of diffusion metrics in the cortex associated with experience-dependent plasticity (3, 4).

## MATERIALS AND METHODS

### Participants

Twenty-one healthy subjects between 18 and 31 years old (11 female; ages: mean ± SD = 23.6 ± 3.1) participated in the study. All subjects were volunteers with no selfreported history of psychiatric, neurological or cognitive impairment. Subjects provided written consent and were compensated for their participation. The experimental procedure was approved by the local Ethics Committee and performed according to the Declaration of Helsinki.

### Experimental Design And Image Acquisition

The dataset used in this study was acquired as part of an ongoing international collaborative project between the Quebec Bio-Imaging Network (QBIN) and the Latin American Brain Mapping Network (LABMAN) aimed at studying plasticity induced by motor learning (9). Participants trained on three different tasks: visuomotor adaptation, motor sequence learning, and a motor control task, which were carried out on three consecutive weeks. Training on each task was separated by a week to minimize interference and carry over effects (10). The order of training on the motor tasks was randomized. Details on the tasks can be found in the Supplementary information.

For each motor task subjects were scanned before training (baseline), as well as 30 minutes and 24 hours after training. Hence, each individual was scanned nine times in total with different magnetic resonance modalities including diffusion-weighted images (DWI) and T1-weighted images.

Magnetic resonance images (MRI) were acquired with a 3T Siemens Tim TRIO scanner using a 12-channel head RF receive coil. Each subject’s head was positioned inside the head coil using the same anatomical landmarks as reference in all sessions. DWI were acquired using a multiband-accelerated sequence (11, 12) using the following parameters: voxel size=2×2×2 mm^3^; field of view =240×240 mm^2^; 30 monopolar gradient directions uniformly distributed (13, 14); 70 axial slices; repetition time=5208 ms; echo time=89 ms; acquisition time=3 minutes and 34 seconds; bandwidth=1488 Hz/Px; multiband acceleration factor=2, SENSE1 *coil-combine* mode, pure axial slice orientation with interleaved slice acquisition, anterior-posterior (A-P) phase encoding direction, with a b-value=1000 s/mm^2^. Eight *b0* volumes (DWI with b-value = 0), were acquired: two at the beginning of the sequence, one at the end and the rest interleaved every five b-1000 volumes (14). Finally, one *b0* volume was acquired with posterior-anterior (P-A) phase encoding direction to correct for susceptibility-induced geometric distortions (15). In addition, one T1-weighted volume was acquired per session using the MP2RAGE WIP sequence (16), with the following parameters: TR = 5000ms; TE = 2.89 ms; flip angle 1 = 4°; flip angle 2=5°; Inversion Time (TI) 1 = 700 ms; TI2 = 2500 ms; BW = 240 Hz/Px; FOV= 256×256 mm^2^; acquisition matrix = 256×256; voxel size = 1×1×1 mm^3^; slices = 176; Parallel acquisition = GRAPPA mode, acceleration factor 2. Acquisition was performed in sagittal slices along the y-z plane of the static magnetic field reference frame.

### Image Pre-Processing

DWI and T1 *DICOM* images were converted into *NIFTI* format using the dcm2nii software (17). Pre-processing steps for DWI were conducted for each of the nine scanning sessions separately using the eddy tool from FSL (version 5.0.9) (18), and included: i) correction of susceptibility-induced distortions (15), ii) correction of eddy currents-induced distortions, head motion correction and b-vector rotation (18). Next, DTIfit was used to fit a diffusion tensor model to generate the scalar measures of interest: *FA* and *MD*. The “halo” of bright voxels that typically surrounds *FA* images due to eddy currents-induced distortions in cerebrospinal fluid (19, 20) was removed by eroding it with a spherical kernel of 6 mm radius (5, 6). A brain mask generated out of the eroded *FA* image was subsequently applied to the associated *MD* and *b0* images. The resulting eroded *FA*, *MD* and *b0* images were then used to evaluate the effect of different normalization approaches. With the ultimate aim of creating an individual T1 template for each subject, pre-processing of the T1 images (T1 uniden) was performed using FreeSurfer’s (21) recon-all pipeline using the default settings.

### Overview Of Normalization Approaches

The DTI scalar maps were normalized using three non-exclusive approaches that varied only in the registration features chosen to bring them into a stereotaxic space. Only DWI data acquired during the baseline and the 24h session of the control task were used to compute the across-session test-retest reproducibility error. From now on we refer to these data as test-retest images. Given the short time interval between these sessions and the absence of a learning manipulation, we assumed that the across-session variability of tissue microstructure metrics would mostly reflect reproducibility errors related to the MRI acquisition protocol and the analysis pipelines.

The following features were assessed in each normalization approach: 1) the registration algorithm used to compute and apply the transformations for image normalization, 2) the registration target, i.e. the image in standard stereotaxic space used as the reference in the normalization process, 3) the use of intermediate targets, i.e., whether DTI images were directly warped to the stereotaxic space or through an intermediate individual template. These normalization approaches were assessed sequentially, taking the feature of the most reproducible pipeline as the default for the subsequent approach. These pipelines with their respective manipulated variables and fixed parameters are outlined in a schematic flowchart (Figure 1), and described in detail as follows.

**Figure 1.**
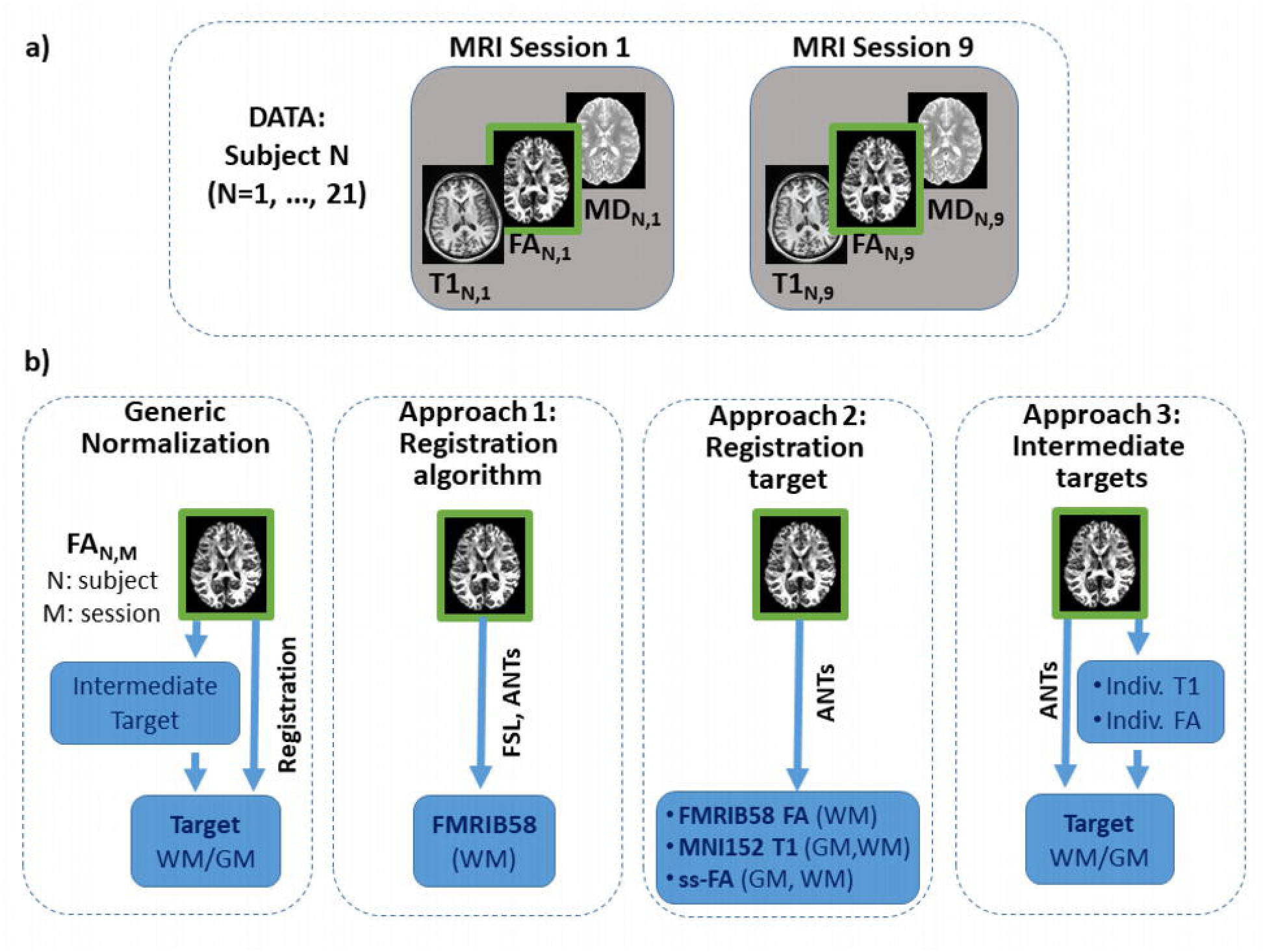
Schematic representation of the data (a) and the various normalization approaches (b). Each subject (N=21) was scanned in multiple sessions (M=9). Each session had a T1 anatomical image and diffusion derived FA and MD maps. The generic normalization approach (b) registers the FA scan of each subject and each session to a target stereotaxic space either directly or through an intermediate subject-specific template derived from the 9 sessions of each subject. Normalization approaches derived from the FA data were applied to the MD data. Three normalization approaches where implemented to assess how various features affected the group test-retest reproducibility errors on WM and GM. Approach 1 manipulated the registration algorithms (FSL and ANTs), using a direct registration to the FMRIB58 atlas that allowed reproducibility estimations only on WM. Approach 2 manipulated the target stereotaxic images (FMRIB58, MNI152 and a study-specific FA (ss-FA)). Approach 3 manipulated the use of an intermediate individual template of the same (Indiv. T1) or different (Indiv. FA) modality of the target atlas (MNI152). Abbreviations: FA, fractional anisotropy; MD, mean diffusivity; WM, white matter; GM, gray matter.

#### Normalization Approach 1: Effect Of Registration Algorithm

Some of the most common pipelines for DTI analyses involve the use of normalization algorithms from FSL (5, 6, 22). However, recently ANTs’ non-linear normalization algorithm (23) has been shown to outperform FSL’s on various metrics (24, 25). To assess the reproducibility of the DTI normalization approach from FSL versus the one from ANTs, we contrasted the following pipelines:

i. A normalization pipeline based on FSL’s registration algorithms.
ii. A normalization pipeline based on ANTs’ registration algorithms.

We used either FSL or ANTs to calculate the linear and non-linear transformations that mapped the *FA* images from each subject’s test and retest sessions to the standard stereotaxic space. These transformations were then concatenated and applied to *FA* and *MD* maps using a single interpolation step.

For linear registration using the FSL’s *FLIRT* tool (FSL version 5.0.9), the default cost function (correlation ratio) was used, which allows the robust registration of all images including those with different contrasts. For non-linear registration using *FNIRT,* the only cost-function presently implemented is the “sum-of-squared differences” (22). We used the configuration file provided in FSL’s toolbox to register the *FA* images to FMRIB58 *FA* template (*FA_2_FMRIB58_1mm.cnf*). Image interpolation was carried out using the tri-linear (default) method.

For linear registration using ANTs (version 2.2.0), translation, rigid and affine transformations were consecutively calculated using the following parameters: Mattes mutual information similarity metric, convergence threshold = 1×10^−6^, convergence window size = 20, gradient step = 0.1. For the non-linear transformation, the symmetric normalization (*SyN*) algorithm (*antsRegistration* command) was used with the following parameters: Mattes mutual information similarity metric, 100×100×50 iterations in three resolution levels with shrink factors = 3×2×1 and smoothing sigmas = 4×2×1, convergence window size = 5, gradient step = 0.2, update field variance in voxel space = 3, total field variance in voxel space = 0. Image interpolation was carried out using the linear method.

Given that FSL’s non-linear registration is optimized to warp images of the same modality (22, 26), *FA* images from the *test* and *retest* sessions were registered to the FMRIB58 *FA* template for both pipelines. Because a large portion of the cortex is missing from the FMRIB58 template, reproducibility was evaluated only for the WM tissue (see Supplementary information for details on the WM mask construction).

#### Normalization Approach 2: Effect Of Registration Target

When one is interested in analyzing diffusion parameters in the neocortex, the template traditionally used in DTI studies (FMRIB58) may be suboptimal (Figure 2). This is due to the fact that the outermost edges of the cortex, containing voxels that are primarily GM or cerebrospinal fluid (CSF) are removed from the template. Here, we proposed the use of the MNI152 T1-based template as the target for registering the DWI images.

**Figure 2.**
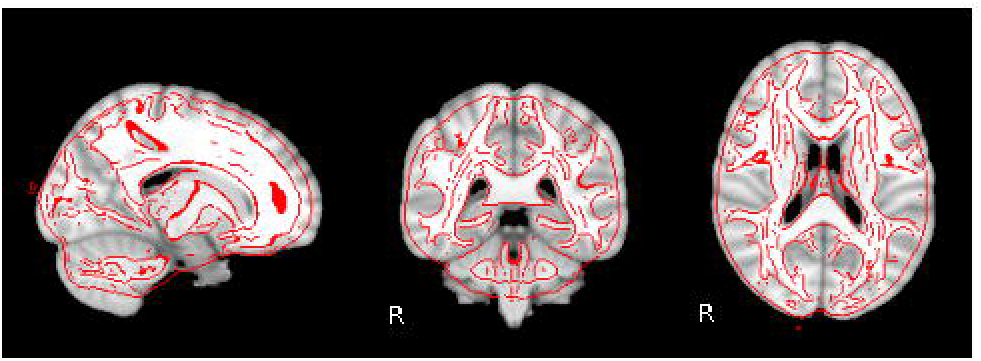
The choice of template in the normalization process matters. Comparison of the two adult human brain templates commonly used as targets for spatial normalization in group studies. The FMRIB58 *FA* template (in red) is shown overlaid on MNI152 T1 template. Note the substantial amount of the cortex excluded from the *FA* template compared to the T1 template.

Given that we are interested in optimizing normalization in the cortex, we also included a study-specific *FA* template (ss-*FA*) generated based on the FA images of all subjects and sessions of our dataset (see the Supplementary information for details on how it was constructed). This template was not thresholded to allow registration at the level of the cortex, which is missing from the FMRIB58. Thus, the reproducibility associated with normalizing *FA* and *MD* images to these three templates was contrasted using the following pipelines:

i. A normalization pipeline using the FMRIB58 *FA* template as the registration target (for white matter reproducibility)
ii. A normalization pipeline using the MNI152 T1 template as the registration target (for gray and white matter reproducibility)
iii. A normalization pipeline using the ss-*FA* template as the registration target (for gray and white matter reproducibility)

*Test* and *retest FA* images were registered to either the FMRIB58 *FA* template, the ss-*FA* template or the MNI152 T1 template by concatenating linear and non-linear transformations using ANTs (the parameters used in the *Registration algorithm* approach were maintained). Transformations calculated for *FA* images were also applied to the corresponding *MD* images. Reproducibility errors in white matter were contrasted for the three templates, whereas reproducibility errors in gray matter were only contrasted for MNI152 T1 and the ss-*FA* templates.

#### Normalization Approach 3: Effect Of Including An Intermediate Target

The first two normalization approaches proposed in this work were based on pipelines in which registration of the *FA* image to the target was performed in a direct fashion. Such direct approach does not differ from what is often used in cros-ssectional studies, but it may not be the best normalization strategy to make the most out of longitudinal data. One of the benefits of having a longitudinal dataset is that within-subject variability may be reduced by taking advantage of the shared structural information of the images acquired at different time points to create an intermediate template for normalization. It has been shown that the use of an intermediate template improves alignment over direct pairwise registration (27–29). Therefore, we evaluated two pipelines using different intermediate templates:

i. A normalization pipeline using an individual *FA* template as intermediate template
ii. A normalization pipeline using an individual T1 template as an intermediate template

The details on how these templates were created are provided in the Supplementary information section. Once the intermediate templates were produced (one individual template per subject), they were registered to the MNI152 T1 using ANTs. Transformations were then applied to the *FA* and *MD* maps in subject space to align them to the MNI152 standard space in a single interpolation step.

To evaluate if adding an intermediate template to the normalization process improved reproducibility in comparison with a direct normalization approach, we contrasted the level of reproducibility from pipelines i) and ii) with that obtained for the pipeline in which normalization was performed directly to the MNI152-T1 template.

### Data Analysis

For all normalization approaches, the performance of the different pipelines was established based on the across-session test-retest reproducibility error using binary masks for each tissue type, namely gray and white matter (refer to the Supplementary information for details on how these masks were created).

#### Across-Session Test-Retest Reproducibility Error

Across-session test-retest reproducibility errors (RE) of *FA* and *MD* were computed in a voxel-wise fashion for each subject and each normalization approach as the absolute difference between the *test* and the *retest* DTI measure, divided by the mean value of both sessions and multiplied by 100 to express it as percent change (8). We chose to use this reproducibility measure because most studies exploring plastic changes in the brain express them as the percentage difference between the follow-up session and the baseline (2–4). Thus, quantifying reproducibility errors as a percentage measure between test and retest sessions would allow us to have a comparable index.

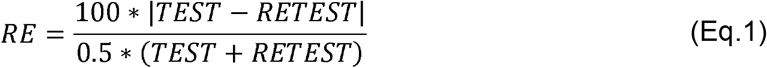

To assess differences in RE for each tissue type, we computed the median RE for each DTI measure over the voxels within a GM and a WM mask. The median was used over the mean because the spatial distribution of REs was right skewed (skewness > 1) for all pipelines.

#### Signal-To-Noise Ratio Assessment

In this study, we focused on the test-retest percent reproducibility error as an index to evaluate a variety of normalization approaches. Yet, a favorable approach in terms of minimizing RE may also impact on the integrity of the image, hindering the signal-to-noise ratio (SNR). To examine if the transformations imposed by the normalization pipelines compromised the integrity of the DTI images, we assessed their impact on the SNR. To obtain the SNR, we computed the mean across each normalized *test* DTI map and divided it by its standard deviation (30). Thus, for each subject we generated one SNR value per DTI measure (one for *MD* and one for *FA*) and tissue type (GM and WM masks). We compared the SNR for the normalization pipelines that overall yielded the lowest reproducibility errors, i.e.:

i. A normalization pipeline registering FA maps directly to MNI152 using ANTs
ii. A normalization pipeline registering FA maps to MNI152 via an intermediate individual FA template using ANTs
iii. A normalization pipeline registering FA maps directly to FMIRB58 using ANTs

#### Statistical Analysis

Statistical analyses were performed using SPSS (IBM SPSS Statistics for Windows, version 25.0). RE and SNR values were statistically compared using a repeated measures analysis of variance (ANOVA), with *pipeline,* DTI measure and tissue type as within-subject factors, when appropriate. Normality of the data was checked using Shapiro-Wilk’s test. For analyses involving comparisons between more than two pipelines, sphericity of the data was tested using Mauchly’s test. In the cases in which the sphericity assumption was not met, a Greenhouse-Geisser degrees of freedom correction was performed. The significance threshold was set at p<0.05. Prior to statistical testing, outliers were detected based on a threshold of three median absolute deviations (MAD) from the group median (31). Using this criterion, one subject was identified as an outlier in the *Registration algorithm* approach and two subjects were considered outliers in the *Intermediate targets* approach. Consequently, they were removed from the corresponding analyses. Post-hoc Tukey tests were used to examine specific differences between pipelines. Bonferroni correction was used to adjust the significance threshold for multiple comparisons.

In the present study, subjects exposed to the control task on the first week of the study were naïve, whereas those who performed this task on the second or third week had been previously exposed to motor learning. In order to rule out potential carry over effects of previous learning on the control DWI scans, we conducted an additional statistical analysis on the 21 participants grouped by the week in which they carried out the control task (refer to the Supplementary information for a description and the results of this analysis).

### Data And Code Availability Statement

The source-code of the pipeline proposed for longitudinal DTI normalization is publicly available in GitHub https://github.com/florjaco/DWIReproducibleNormalization. The dataset used for this work is also available upon request.

## RESULTS

### Registration Algorithm

The quantitative assessment of the RE for both algorithms is shown in Figure 3, Supplementary Figure 1 and Supplementary Table 1. The upper row shows the distributions of reproducibility errors in white matter for *MD* (left) and *FA* (right), respectively. For both metrics we found that ANTs yielded lower percent reproducibility errors than FSL: *MD* = Mean±SE: RE=4.61±0.09% for FSL vs. RE=4.22±0.08% for ANTs (F(1,19)=492.164, p<0.0001), *FA* = Mean±SE: RE=8.44±0.11% for FSL vs RE=6.04±0.07% for ANTs (F(1,19)= 1930.55, p<0.0001). The bottom row of Figure 3 shows the color coded voxel-wise spatial map of the mean RE computed across the WM mask. Note that the reproducibility errors were higher for *FA* than for *MD* in both normalization pipelines (main effect of DTI measure F(1,19)=3465.33, p<0.0001, with mean RE=7.24±0.09% for *FA* vs RE=4.42±0.08% for *MD*).

**Figure 3.**
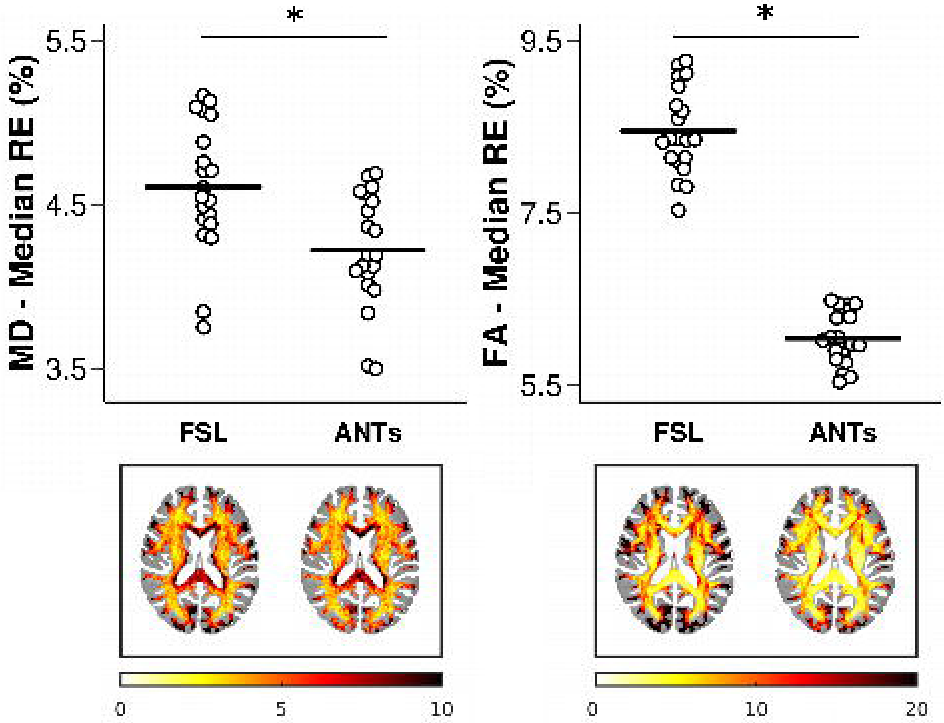
The choice of non-linear registration algorithm (FSL or ANTs) used with the FMRIB58 *FA* template affects test-retest reproducibility of diffusion scalars in white matter. ANTs yielded significantly lower reproducibility errors than FSL (*p<0.0001). The upper plots show the median percent test-retest reproducibility error for *MD* (left) and *FA* (right). Note that different scales are used for panels showing *MD* and *FA* results. The bottom row shows the color-coded anatomical distribution of percent reproducibility errors in the white matter mask of the *FA* template; axial slice coordinate: z=19 mm. In order to show more clearly the different dynamic ranges in the voxel-wise colormaps, we used a different range of values for reproducibility errors in *MD* and *FA*. Acronyms: mean diffusivity (*MD*), fractional anisotropy (*FA*), test-retest reproducibility error (RE).

### Registration Target

The quantitative assessment of the reproducibility error associated with the target used for registration is shown in Figure 4, Supplementary Figure 2 and Supplementary Tables 2 and 3. Reproducibility errors differed in WM across the three pipelines used to test the effect of the registration target: MNI152, FMRIB58 and ss-*FA* (F(1.25, 25.05)=116.51, p<0.0001 for FA and F(1.31, 26.42)=127.93, p<0.0001 for MD). Specifically, the MNI152 was associated with lower reproducibility errors for MD (p<0.0001), whereas the FMRIB58 was associated with lower reproducibility errors for *FA* (p<0.0001). In gray matter, RE was significantly different for *FA* (F(1,20)=146.75, p<0.0001) but not for *MD* (F(1,20)=2.05, p=0.168). Specifically, the MNI152 yielded a lower RE than the ss-*FA* template for *FA* (p<0.0001) but a similar RE for MD.

**Figure 4.**
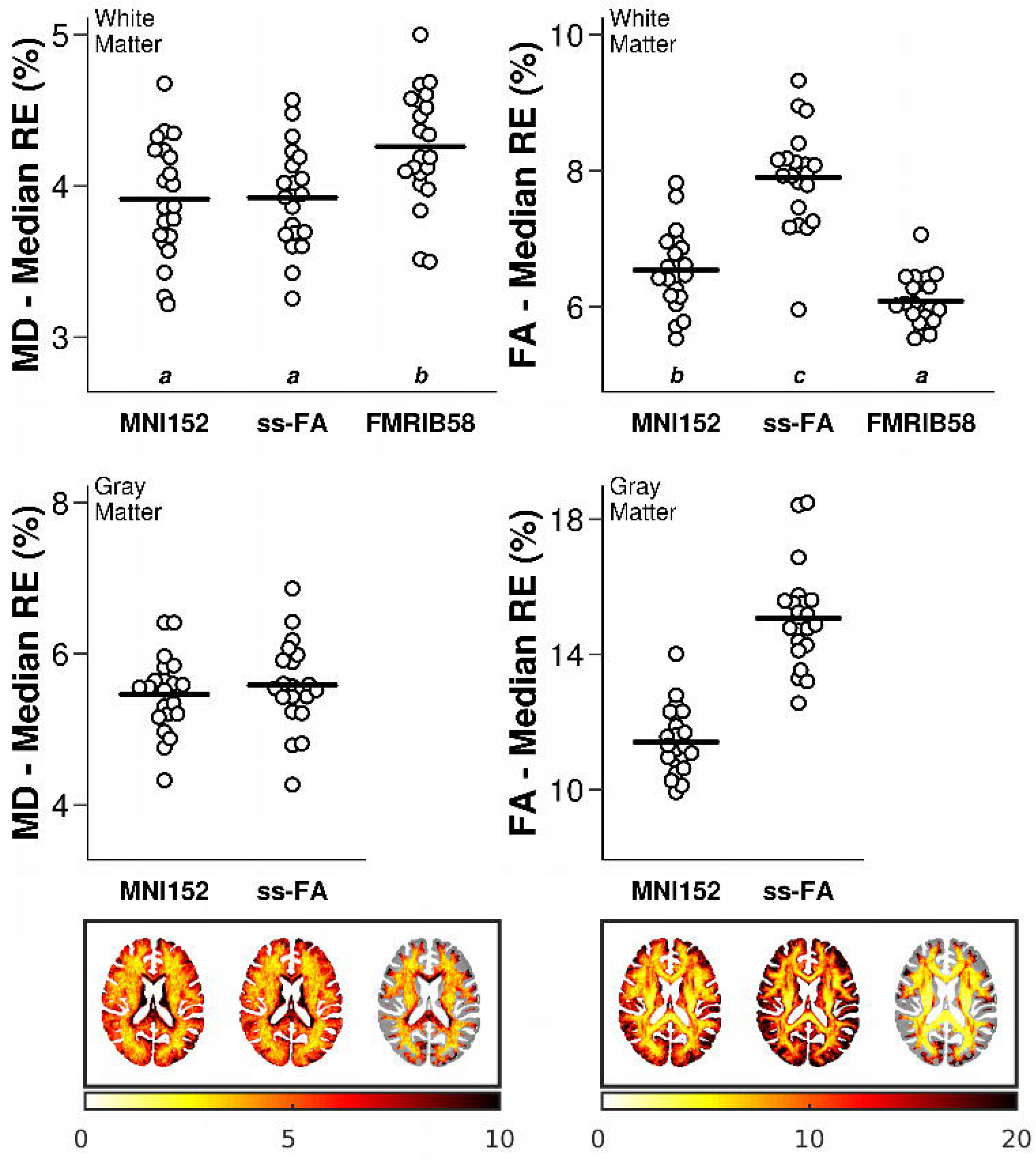
The choice of registration target affects reproducibility errors in gray and white matter. Shown are the median RE (plots), and the color-coded anatomical distribution of RE for each diffusion metric (*MD* and *FA*) and each target template (MNI152 T1, ss-*FA* and FMRIB58 *FA*). As indicated in Eq. (1) RE is expressed as percent change. Note that different scales are used for panels showing *MD* and *FA* results. In order to show more clearly the different dynamic ranges in the voxel-wise colormaps, we used a different range of values for reproducibility errors in *MD* and *FA*. Letters above the horizontal axis represent the compact display of all pair-wise comparisons using Tukey’s test. Different letters express differences between pipelines with an adjusted p-value<0.05. Same letters indicate no statistical differences. Acronyms: mean diffusivity (*MD*), fractional anisotropy (*FA*), test-retest reproducibility error (RE).

In line with the previous normalization approach, RE was higher for *FA* than *MD* for all the pipelines considered (main effect of DTI measure in WM F(1,20)=1469.51, p<0.0001, with mean RE=6.84±0.10% for *FA* vs RE=4.03±0.79% for *MD;* main effect of DTI measure in GM F(1,20)=1926.82, p<0.0001, with mean RE=13.25±0.23% for *FA* vs RE=5.52±0.11% for *MD*). Reproducibility errors were higher in GM than in WM, regardless of pipeline (ss-*FA* and MNI152) and diffusion metric (main effect of tissue type F(1,20)=1602.66, p<0.0001, with mean RE=9.39±0.16% for GM vs RE=5.57±0.09% for WM). Voxel-wise distribution of RE is depicted at the bottom of Figure 4.

### Use Of Intermediate Targets

The quantitative assessment of the reproducibility error associated with normalization via an intermediate target is shown in Figure 5, Supplementary Figure 3 and Supplementary Tables 4 and 5. The reproducibility error in white matter differed significantly both for *MD* (F(2,36)=7.27; p=0.002) and *FA* (F(2,36)=72.91; p<0.0001). A post-hoc test showed that using an individual *FA* template as an intermediate step in the normalization process yielded the lowest RE for *FA* in white matter (Figure 5, top panel). Note that the RE for this pipeline did not differ from that obtained using the FMRIB58 template (F(1,20)<0.001; p=0.99; see the inset of Figure 5 top right panel), suggesting that both pipelines perform similarly for this measure. In contrast, normalization via the T1-based individual template produced higher RE than all of the other pipelines (p<0.004), except for the MNI152 (for MD in WM). The reproducibility error in gray matter also differed significantly for both *MD* (F(2,36)=22.17, p<0.0001) and *FA* (F(2,36)=101.60, p<0.0001). Yet, this difference was accounted for the T1-based individual template, which yielded the highest RE both for FA and MD (p<0.003). Voxel-wise maps for the mean RE across subjects are shown in Figure 5 to illustrate the RE distribution.

**Figure 5.**
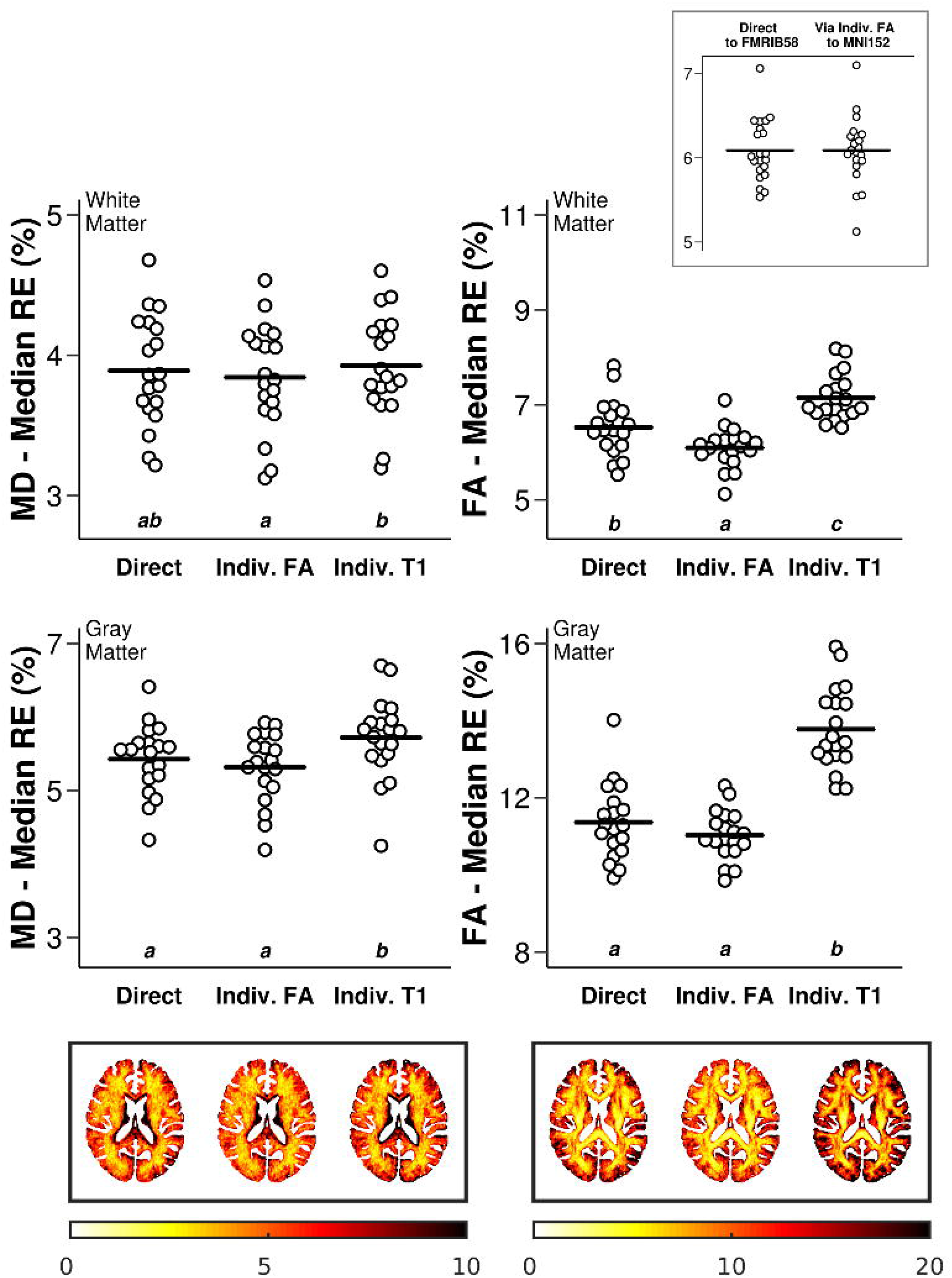
The choice of an intermediate individual template affects reproducibility error in GM and WM. Shown are the median RE (plots), and the color-coded anatomical distribution of RE for *MD* (left) and *FA* (right) for white and gray matter. As indicated in Eq. (1) RE is expressed as percent change. **Direct**: direct normalization, **Indiv. FA**: normalization via an individual *FA* template, and **Indiv. T1**: normalization via an individual T1 template. The inset shows that the pipeline that performs best for FA in white matter (Indiv. *FA*) yields comparable REs when registering directly to the FMRIB58. Note that different scales are used for panels showing *MD* and *FA* results. Letters above the horizontal axis represent the compact display of all pairwise comparisons using Tukey’s test. Different letters express differences between pipelines with an adjusted p-value<0.05. Same letters indicate no statistical differences. Acronyms: mean diffusivity (*MD*), fractional anisotropy (*FA*), test-retest reproducibility error (RE), gray matter (GM), white matter (WM).

As for previous normalization approaches, the reproducibility error associated with *FA* was higher than for *MD* for all pipelines and across tissues (main effect of DTI measure F(1,18)=2172.03, p<0.0001, with mean RE=9.33±0.13% for *FA* vs RE=4.69±0.09% for *MD*). Also note that reproducibility errors were higher in GM than in WM, regardless of the pipeline and diffusion metric (main effect of tissue type F(1,18)=2850.41, p<0.0001, with mean RE=8.77±0.12% for GM vs RE=5.24±0.09% for WM).

### Signal-To-Noise Ratio Assessment

The quantitative assessment of the SNR associated with the normalization pipelines that overall yielded the lowest reproducibility errors is shown in Figure 6. Statistical assessment of SNR in white matter identified a significant effect of pipeline for *MD* (F(1.40,27.94)=79.59, p<0.0001) and *FA* (F(2,40)=1205.99, p<0.0001). A post-hoc test conducted on *MD* revealed that the use of an individual *FA* template as an intermediate step for normalization yielded the highest SNR (p<0.01). On the other hand, the direct normalization of FA images to the MNI152 template yielded the highest SNR for FA (p<0.0001). Of note, direct normalization to the FMRIB58 FA template yielded the lowest SNR for both measures (p<0.01).

**Figure 6.**
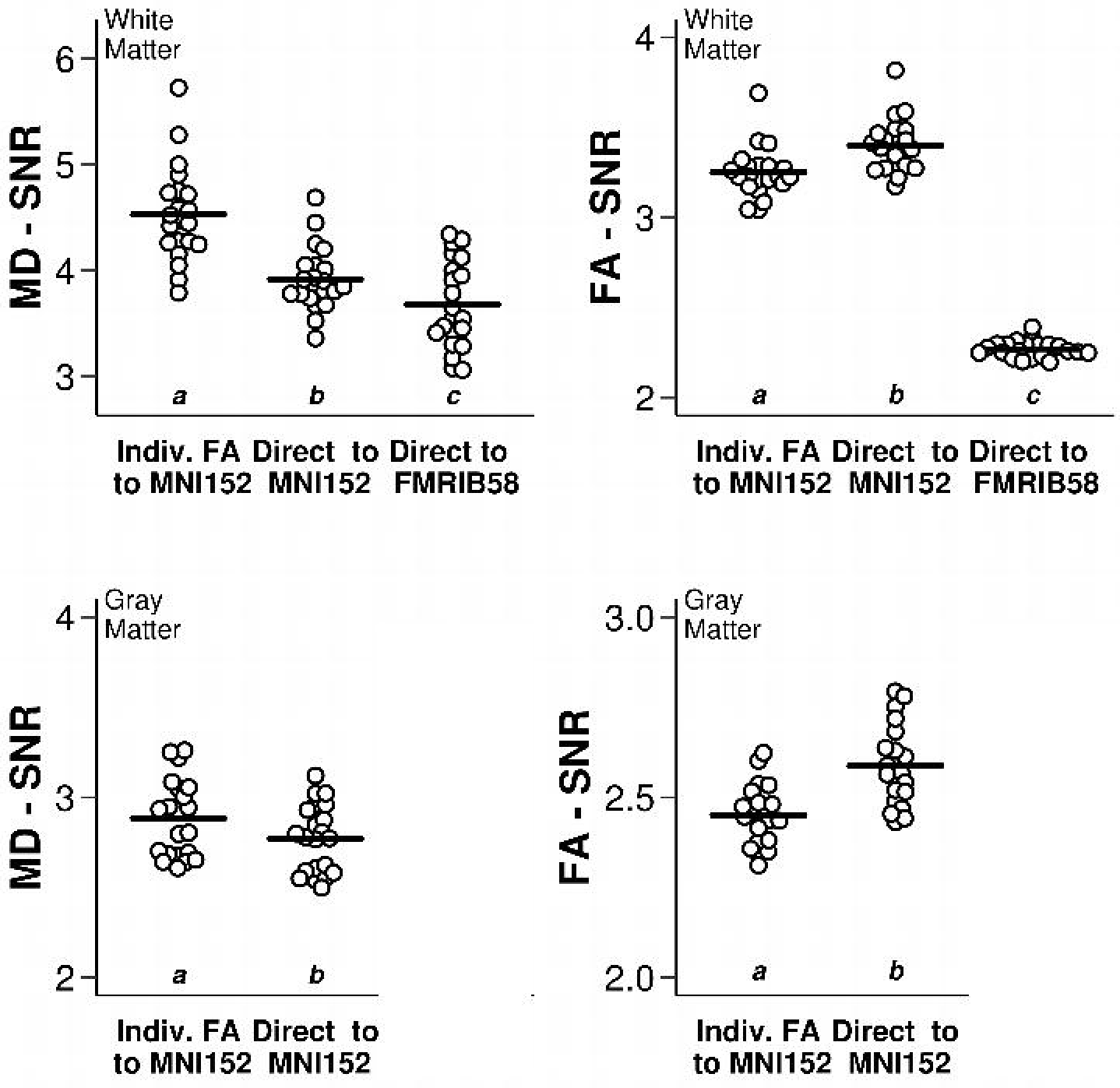
The use of an intermediate FA template for brain spatial normalization affects the global signal-to-noise ratio (SNR) in *MD* and *FA* maps. Shown are the SNR for MD (left) and FA (right) for white and gray matter. **Direct to MNI152**: direct normalization of FA maps to MNI152, **Indiv. FA to MNI152**: normalization of FA maps to MNI152 via an individual FA template, and **Direct to FMRIB58**: direct normalization of FA maps to FMRIB58. Note that different scales are used for all panels. Letters above the horizontal axis represent the compact display of all pairwise comparisons using Tukey’s test. Different letters express differences between pipelines with an adjusted p-value<0.05. Same letters indicate no statistical differences. Acronyms: mean diffusivity (*MD*), fractional anisotropy (*FA*).

Statistical assessment of SNR in gray matter also identified a significant effect of pipeline both for *MD* (F(1,20)=57.45; p<0.0001) and *FA* (F(1,20)=100.60; p<0.0001). Normalization using an intermediate *FA* template yielded the highest SNR (p<0.0001) on MD. On the other hand, normalizing FA images directly to the MNI152 template produced the highest SNR (p<0.0001) on FA. Refer to Supplementary Tables 6 and 7 for additional information regarding differences between these pipelines and post-hoc comparisons.

## DISCUSSION

Brain diffusion MRI scalar maps like fractional anisotropy and mean diffusivity provide an increasingly valuable non-invasive tool to quantify structural plasticity (1, 2, 4, 32). Yet, longitudinal changes in gray matter are usually overlooked in traditional pipelines of DTI analysis, generally focused on white matter tissue. Here, we sought to assess spatial normalization approaches that would enhance the probability to detect longitudinal changes in diffusion scalar maps at the level of the cortex. Previous studies have evaluated the reproducibility of different aspects of the normalization process such as the registration algorithm or the use of an individual template as an intermediate step (32–34). However, these attempts usually assessed one or two registration traits at a time and they put emphasis on reproducibility within white matter. Here, we explored the impact of varying different features of the spatial normalization process on the across-session test-retest reproducibility error for both gray matter and white matter. We did so by systematically and sequentially comparing the test-retest reproducibility from different normalization approaches. We first evaluated the effect of non-linear registration algorithm (ANTs and FSL). We then examined the choice of registration target (FMRIB58-FA, MNI152-T1 and a study-specific FA template). Finally we evaluated the effect of including an intermediate individual template (T1-based or FA-based) prior to the registration to the stereotaxic space.

We found that the ANTs registration algorithm yields the best reproducibility in white matter. This finding is in line with previous work showing that ANTs is associated with a low rate of false positives (25) and good alignment of white matter tracts (35). Moreover, we found that when using ANTs as the registration algorithm, the choice of the target image affected the test-retest reproducibility of diffusion metrics in white matter, in a way that depended on the diffusion scalar of interest (FA or MD). Specifically, using the FMRIB58 template as the target of normalization, improved reproducibility for FA over that achieved with the MNI152, whereas the use of the MNI152 outperformed the FMRIB58 for MD. These two registration targets cannot be directly compared in gray matter due to their unequal cortical coverage. To overcome this limitation, we created a study-specific *FA* template (ss-*FA*) which, unlike FMRIB58, was not thresholded. We found that using ANTs as the registration algorithm and MNI152 as target improved reproducibility both for FA and MD in gray matter. Therefore, it appears that the MNI152 outperforms FA-based templates for all DTI measures except for FA in white matter.

Yet, when interested in detecting microstructural changes in the whole brain, it is desirable to use a single pipeline to optimize registration for both DTI measures and tissue types. To this aim, we decided to include an intermediate individual template in the registration of DTI to MNI152 to improve further the registration of FA and MD in white matter. This procedure has been shown to enhance registration in longitudinal datasets (7). Accordingly, we found that the use of an individual FA template produced the highest reproducibility for FA in white matter, whereas it did not differ from directly registering to the MNI152 in gray matter. It is important to note that normalization via an individual FA template yielded a comparable level of RE to that achieved when using ANTs for registration to the FMRIB58 template. Therefore, even though using an individual FA template does not improve registration further in the cortex, it provides a unifying pipeline that is optimal for both DTI metrics and tissue types. The fact that this pipeline also shows higher SNR than the FMRIB58 pipeline strengthens this result. In sum, our findings suggest that registration via an individual template is beneficial when its modality matches that of the moving image (i.e., the image to be warped).

Unexpectedly, using a T1-based individual template, created with FreeSurfer’s longitudinal pipeline (27), did not improve reproducibility for any DTI measures or tissue types. This finding was unpredicted given the high tissue contrast and spatial resolution of these images, and the fact that the non-linear transformation was performed within-modality (T1 to MNI152 T1 template). We chose FreeSurfer to create the individual T1-based templates over the *antsMultivariateTemplateConstruction* tool (used to construct the individual FA-based templates) because it is currently the state-of-the-art pipeline for unbiased T1 longitudinal analysis and template creation (27). Yet, it is important to remark that FreeSurfer’s approach for template creation is surface-based, while normalization of this template to MNI152 with ANTs was performed using volumetric transformations. This may have rendered overall registration suboptimal.

One systematic bias we observed across all normalization approaches was the magnitude of whole-brain reproducibility errors, which was higher, almost twofold, for *FA* than for *MD,* regardless of the feature examined and type of tissue. These findings are in agreement with previous studies, showing that *FA* has a higher spatial variability of reproducibility errors across brain regions than *MD* (30, 33, 36). Moreover, both *FA* and *MD* were more reproducible in white matter than in gray matter. This tissue-dependent difference in reproducibility errors has been reported previously for *FA* in white and subcortical gray matter (30, 37) and for both *FA* and *MD* in gray and white matter (36). Cortical voxels can be affected by partial-volume effects in the GM/CSF interface, which could explain why areas of gray matter in the cortex show higher reproducibility errors (38)(for example, see voxel-wise maps of RE in Figure 5). To reduce partial volume effects in our study, test-retest reproducibility errors were computed over data that was normalized to a standard stereotaxic template using a single interpolation step to avoid multiple resampling and the associated degradation of the images. In addition, CSF was removed and no spatial smoothing was applied. Although these precautions cannot rule out the occurrence of partial volume effects, the fact that reproducibility error was computed on scans acquired within a short (24 h) time frame, suggests that the extent of partial volume effects were comparable across sessions. Therefore, our results likely reflect differences in the impact of the normalization features rather than artifacts driven by differences in partial volume.

Despite its strengths, our work presents limitations. First, the reproducibility error was assessed at the level of tissue by computing the median of voxel-wise test-retest measures of reproducibility within whole-brain white matter and gray matter regions of interest. Although this approach facilitates extracting conclusions on the global impact of the contrasted features, it overlooks any spatial variations. Statistical analysis over the raw voxel-wise test-retest reproducibility error maps would allow estimating the influence of other factors such as anatomical variability, signal-to-noise ratio and image inhomogeneities. Second, the reproducibility error provides only one way to measure the reliability of a spatial normalization approach. Other aspects such as the specificity and sensitivity of the method in question would allow establishing the false positive and negative rates, and the minimum detectable difference of the DTI measure of interest. The specificity and sensitivity of different normalization approaches have been addressed for *FA* maps in white matter (25, 32, 39), and may be extended to other DTI measures. Finally, although our work suggests that normalizing DTI images to the MNI152 via an individual FA template optimized registration for white and gray matter, other combination of features may improve it even further, or make it more amenable for patient populations. For example, the use of full-tensor information for the creation of the within-subject template and for normalization could be a possible alternative to further improve reproducibility (32, 40). Future studies are needed to address these possibilities.

In conclusion, our work explored different normalization approaches with the aim of improving the detection of subtle longitudinal changes in microstructure, both in gray and white matter tissue. We showed that a normalization approach using ANTs to register *FA* images to the MNI152 via an individual *FA* template minimized reproducibility errors for *FA* and *MD*. The performance of this pipeline was comparable to traditional algorithms currently used to assess variations in microstructure in white matter tracts, while allowing quantification of diffusion properties in gray matter, thereby opening a window to assess plasticity at the cortical level.

## ACKNOWLEDGMENTS

We want to thank Chiara Maffei for her advice in early phases of this work, Lisa Novello for her help testing the distributed analysis scripts and related documentation, and Arnaud Boré for technical assistance. We also would like to thank the University of Buenos Aires and, especially, Vice-rector Juan Pablo Mas Velez for their support in getting the first MRI facility dedicated to research going in Buenos Aires.

