## Supplementary information for "Improving spatial normalization of brain diffusion MRI to measure longitudinal changes of tissue microstructure in the cortex and white matter"

**METHODS**

***Motor Learning Tasks***

The MRI dataset used in this study was acquired as part of an ongoing international collaborative project between the Quebec Bio-Imaging Network (QBIN) and the Latin American Brain Mapping Network (LABMAN) aimed at studying plasticity induced by learning (1). Participants trained on three different tasks: visuomotor adaptation (VMA), motor sequence learning (MSL), and a motor control task, which were carried out in three consecutive weeks. In the visuomotor adaptation task, subjects were required to move a cursor from a starting point in the center of a screen to different visual targets using a joystick controlled with the index and thumb of their right hand. The task was carried out in the presence of a constant visual rotation of 40° that altered the movement of the cursor. Throughout adaptation subjects learned to adjust the trajectory of the cursor gradually until it reached the target correctly (2). The motor sequence learning task required subjects to press a series of four keys with the fingers of the left hand following a sequence of movements of five elements, which were previously explicitly memorized (3, 4). The objective of the task was to execute the sequence as quickly and accurately as possible. The control task consisted of a go/no-go paradigm in which subjects had to press a button with their right index finger in response to a green target appearing on a screen (p=0.9) and withhold from responding whenever the target was red (p=0.1).

***Individual Template Construction***

It has previously been shown that the use of an intermediate template improves alignment over direct pairwise registration (5–7). Therefore, we evaluated two pipelines using different intermediate templates:

i. A normalization pipeline using an individual *FA* template as intermediate template
ii. A normalization pipeline using an individual T1 template as intermediate template

When having only two time points for the same subject, a common strategy to create an individual template is to refer both images to their halfway point or mid-space (8). In contrast, when the number of time points is larger, as in this case in which 9 images were obtained per subject, creating a mid-space for all time points is cumbersome. Given that the complete longitudinal data set included nine time-points per subject, we included all nine images to obtain the best representation of each subject’s brain when generating the individual template. This is in line with previous work in which subject-specific templates of other modalities were generated using all time-points as input for template creation (6, 9). The individual FA template was created using *antsMultivariateTemplateConstruction* tool with the following parameters: rigid-body registration of inputs for the creation of an initial template used as seed, gradient step size=0.2, cross-correlation similarity metric, Greedy-SyN transformation model used for registration. The individual T1 template was created using FreeSurfer’s longitudinal stream (6). Specifically, an unbiased within-subject template space and image (10) is created using robust, inverse consistent registration (11).

For each subject, both the individual FA template and the individual T1 template were registered to MNI152 T1 template using ANTs. The parameters used for this step were the same as the ones described in the *Registration Algorithm* section. The template construction tools automatically provide the transformations that map each one of the input images to the output template. Hence, for the normalization of images in pipeline i), the transformations that map the *test* and *retest* *FA* images to the individual template were concatenated to those that map the individual template to MNI152 space. For pipeline ii), the *test* and *retest* *FA* images were first linearly registered to their corresponding T1 image (via *b0*) using ANTs. The transformations that aligned the *FA* images to the T1, the T1 to the individual T1 template, and the individual T1 template to MNI152 space were concatenated. Transformations were applied to the *FA* and *MD* maps in subject space to align them to the MNI152 standard space in a single interpolation step.

***Study-Specific FA Template Construction***

For the creation of the study-specific *FA* (ss-*FA*) template, individual *FA* templates from all subjects (twenty-one) were used as input to the *antsMultivariateTemplateConstruction* tool (12, 13). The parameters were the same as the ones described for the construction of the individual *FA* templates. The ss-*FA* template was then non-linearly registered to MNI152 space using ANTs. The transformations that map the *test* and *retest* *FA* images to the individual template, the corresponding individual template to the ss-*FA* template and the ss-*FA* template to MNI152 space were concatenated. Then, these transformations were applied to the *FA* and *MD* maps in subject space to align them to standard space in a single interpolation step.

***Creation Of White And Gray Matter Masks***

Masks were created by segmenting the MNI152 T1 template into three tissues (GM, WM and CSF) using FSL’s FAST tool (14). The GM component obtained from the segmentation was used to create an atlas GM mask. An additional subcortical mask was created using the Harvard-Oxford subcortical structural atlas (15) (thresholded=60% probability) to include a series of structures of interest that were missing from the automatically generated GM component. This included the right and left pallidum, putamen, caudate and thalamus. Cortical and subcortical GM masks were combined and binarized into a unique GM mask. On the other hand, the WM mask was created based on the intersection of the components automatically generated by segmenting WM tissues from the MNI152 template and the FMRIB58 template (the FMRIB58 is substantially smaller due to thresholding). This allowed comparing RE in WM for images normalized with either template.

**RESULTS**

***Test-Retest Reproducibility Error***

In this section, we present additional statistics that complement the data presented in the Results section of the manuscript. We show the descriptive statistics and the results of the repeated-measures ANOVA performed on the test-retest reproducibility error for each normalization approach i.e., *Registration algorithm, Registration target and Intermediate targets* for FA (fractional anisotropy) and MD (mean diffusivity) in WM and GM, where appropriate. We also present the results of post-hoc Tukey tests on these measures.

**Supplementary Table 1:** Test-retest across-session reproducibility error (RE, mean ± standard error) associated with the *Registration algorithm* normalization approach (FSL vs ANTs).

| *Reproducibility error (Mean±Standard Error %) in white matter* | | | |
| --- | --- | --- | --- |
|  | **FSL** | **ANTs** | Repeated Measures ANOVA results |
| *RE of MD* | 4.61±0.09 | 4.22±0.08 | F(1,19)=492.164, p<0.0001 |
| *RE of FA* | 8.44±0.11 | 6.04±0.07 | F(1,19)=1930.55, p<0.0001 |

**Supplementary Table 2:** Test-retest across-session reproducibility error (RE) associated with the *Registration target* normalization approach (FMRIB58, MNI152 or a study-specific group template – ss-*FA*).

| *Reproducibility error (Mean±Standard Error %) in white matter* | | | | |
| --- | --- | --- | --- | --- |
|  | **FMRIB58** | **MNI152** | **ss-*FA*** | Repeated Measures ANOVA results |
| *RE of MD* | 4.26±0.08 | 3.92±0.08 | 3.92±0.07 | F(1.32,26.42)=127.93; p<0.0001 |
| *RE of FA* | 6.09±0.08 | 6.54±0.13 | 7.90±0.16 | F(1.25,25.05)=116.51; p<0.0001 |
| *Reproducibility error (Mean±Standard Error %) in gray matter* | | | | |
|  |  | **MNI152** | **ss-*FA*** | Repeated Measures ANOVA results |
| *RE of MD* |  | 5.46±0.11 | 5.59±0.13 | F(1,20)=2.05; p=0.168 |
| *RE of FA* |  | 11.42±0.22 | 15.08±0.33 | F(1,20)=146.75; p<0.0001 |

**Supplementary Table 3:** Post-hoc Tukey test on the reproducibility error associated with the Registration t*arget* normalization approach in white matter (FMRIB58, MNI152 or a study-specific group template – ss-*FA*). Differences between means are presented as mean ± standard error.

| *Differences of reproducibility error of MD in white matter*  *(Mean±Standard Error %)* | | | |
| --- | --- | --- | --- |
| *Pipeline i* | *Pipeline j* | *Difference between means (i-j)* | *Significance^#^* |
| **ss-*FA*** | **MNI152** | 0.009±0.026 | 1 |
|  | **FMRIB58** | -0.335±0.030 | <0.0001 |
| **MNI152** | **FMRIB58** | -0.344±0.014 | <0.0001 |
| *Differences of reproducibility error of FA in white matter*  *(Mean±Standard Error %)* | | | |
| *Pipeline i* | *Pipeline j* | *Difference between means (i-j)* | *Significance^#^* |
| **ss-*FA*** | **MNI152** | 1.356±0.143 | <0.0001 |
|  | **FMRIB58** | 1.810±0.147 | <0.0001 |
| **MNI152** | **FMRIB58** | 0.454±0.059 | <0.0001 |

^#^Significance: adjusted p-values with Bonferroni correction are shown (i.e., p-value multiplied by nc, the number of comparisons). A p-value=1 means the unadjusted p-value is greater than or equal to 1/nc.

**Supplementary Table 4:** Test-retest across-session reproducibility error (RE) associated with the *Intermediate targets* normalization approach: Direct, via an individual T1 template (Indiv. T1), or via an individual FA template (Indiv. FA).

| *Reproducibility error (Mean±Standard Error %) in white matter* | | | | |
| --- | --- | --- | --- | --- |
|  | **Direct** | **Indiv. T1** | **Indiv. FA** | rm-ANOVA results |
| *RE of MD* | 3.89±0.09 | 3.92±0.09 | 3.84±0.09 | F(2,36)=7.27, p=0.002 |
| *RE of FA* | 6.53±0.13 | 7.15±0.11 | 6.10±0.10 | F(2,36)=72.91, p<0.0001 |
| *Reproducibility error (Mean±Standard Error %) in gray matter* | | | | |
|  | **Direct** | **Indiv. T1** | **Indiv. FA** | rm-ANOVA results |
| *RE of MD* | 5.43±0.11 | 5.72±0.13 | 5.32±0.11 | F(2,36)=22.17, p<0.0001 |
| *RE of FA* | 11.37±0.23 | 13.78±0.25 | 11.03±0.15 | F(2,36)=101.60, p<0.0001 |

**Supplementary Table 5:** Post-hoc Tukey test on the reproducibility error associated with the *Intermediate targets* normalization approach: Direct, via an individual T1 template (Indiv. T1) or via an individual FA template (Indiv. FA).

| *Differences of reproducibility error of MD in white matter*  *(Mean±Standard Error %)* | | | |
| --- | --- | --- | --- |
| *Pipeline i* | *Pipeline j* | *Difference between means (i-j)* | *Significance^#^* |
| **Indiv. FA** | **Indiv. T1** | -0.082±0.022 | 0.004 |
|  | **Direct** | -0.046±0.020 | 0.101 |
| **Indiv. T1** | **Direct** | 0.036±0.023 | 0.399 |
| *Differences of reproducibility error of FA in white matter*  *(Mean±Standard Error %)* | | | |
| *Pipeline i* | *Pipeline j* | *Difference between means (i-j)* | *Significance^#^* |
| **Indiv. FA** | **Indiv. T1** | -1.059±0.087 | <0.0001 |
|  | **Direct** | -0.430±0.071 | <0.0001 |
| **Indiv. T1** | **Direct** | 0.629±0.103 | <0.0001 |
| *Differences of reproducibility error of MD in gray matter*  *(Mean±Standard Error %)* | | | |
| *Pipeline i* | *Pipeline j* | *Difference between means (i-j)* | *Significance^#^* |
| **Indiv. FA** | **Indiv. T1** | -0.401±0.061 | <0.0001 |
|  | **Direct** | -0.111±0.050 | 0.118 |
| **Indiv. T1** | **Direct** | 0.290±0.073 | 0.003 |
| *Differences of reproducibility error of FA in gray matter*  *(Mean±Standard Error %)* | | | |
| *Pipeline i* | *Pipeline j* | *Difference between means (i-j)* | *Significance^#^* |
| **Indiv. FA** | **Indiv. T1** | -2.747±0.226 | <0.0001 |
|  | **Direct** | -0.337±0.161 | 0.153 |
| **Indiv. T1** | **Direct** | 2.410±0.236 | <0.0001 |

^#^Significance: adjusted p-values with Bonferroni correction are shown (i.e., p-value multiplied by nc, the number of comparisons). A p-value=1 means the unadjusted p-value is greater than or equal to 1/nc.

***Signal-To-Noise Ratio Assessment***

Below, we present the descriptive statistics and the results of the repeated-measures ANOVA performed on the signal-to-noise ratio (SNR) for FA and MD in white and gray matter. We also present the results of post-hoc Tukey tests on these measures.

**Supplementary Table 6:** Signal-to-noise ratio (SNR) associated with different normalization pipelines: Direct to FMRIB58, Direct to MNI152, via an individual FA template (Indiv. FA) to MNI152.

| *Signal-to-noise ratio (Mean±Standard Error) in white matter* | | | | |
| --- | --- | --- | --- | --- |
|  | **Direct to FMRIB58** | **Direct to MNI152** | **via Indiv. FA to MNI152** | rm-ANOVA results |
| *SNR of MD* | 3.68±0.09 | 3.92±0.07 | 4.53±0.10 | F(1.40,27.94)=79.59, p<0.0001 |
| *SNR of FA* | 2.30±0.01 | 3.40±0.03 | 3.25±0.03 | F(2,40)=1205.99, p<0.0001 |
| *Signal-to-noise ratio (Mean±Standard Error) in gray matter* | | | | |
|  |  | **Direct to MNI152** | **via Indiv. FA to MNI152** | rm-ANOVA results |
| *SNR of MD* |  | 2.77±0.04 | 2.88±0.05 | F(1,20)=57.45, p<0.0001 |
| *SNR of FA* |  | 2.59±0.02 | 2.45±0.02 | F(1,20)=100.60, p<0.0001 |

**Supplementary Table 7:** Post-hoc Tukey test on the signal-to-noise ratio associated with different normalization pipelines: Direct to FMRIB58, Direct to MNI152, via an individual FA template (Indiv. FA) to MNI152.

| *Differences of signal-to-noise ratio (Mean±Standard Error) of MD in white matter* | | | |
| --- | --- | --- | --- |
| *Pipeline i* | *Pipeline j* | *Difference between means (i-j)* | *Significance^#^* |
| **via Indiv. FA to MNI152** | **Direct to MNI152** | 0.610±0.044 | <0.0001 |
|  | **Direct to FMRIB58** | 0.850±0.086 | <0.0001 |
| **Direct to MNI152** | **Direct to FMRIB58** | 0.240±0.072 | 0.009 |
| *Differences of signal-to-noise ratio (Mean±Standard Error) of FA in white matter* | | | |
| *Pipeline i* | *Pipeline j* | *Difference between means (i-j)* | *Significance^#^* |
| **via Indiv. FA to MNI152** | **Direct to MNI152** | -0.149±0.024 | <0.0001 |
|  | **Direct to FMRIB58** | 0.981±0.024 | <0.0001 |
| **Direct to MNI152** | **Direct to FMRIB58** | 1.130±0.027 | <0.0001 |

^#^Significance: adjusted p-values with Bonferroni correction are shown (i.e., p-value multiplied by nc, the number of comparisons). A p-value=1 means the unadjusted p-value is greater than or equal to 1/nc.

***Task Order Does Not Influence Test-Retest Reproducibility Error***

To explore whether the control condition (*go-nogo* task) used for the test-retest reproducibility assessment was biased by previous learning, we grouped the 21 participants according to the week of the experiment in which they carried out the control task (n=7 subjects/group): week1, week2 and week3. Week1 group included those subjects that carried out the control task on the first week of the experiment. Week2 group included those subjects that were first exposed to a motor learning task (either MSL or VMA) and carried out the control task on the second week of the experiment. Week3 group, included those subjects that were exposed to the control task after both MSL and VMA, during the third week of the experiment. For each normalization approach (*Registration algorithm*, *Registration target* and *Intermediate targets*), DTI measure (MD and FA), and tissue type (white and gray matter) we ran a two-way mixed ANOVA with “Pipeline” as within-subjects measure and “Group (Week1 through 3)” as between-subjects measure. We found no significant interaction between Pipeline and Group, suggesting that reproducibility errors were similar across groups for the normalization features under study. Moreover, there was no significant effect of Group on reproducibility error, but a significant effect of Pipeline (stats are reported below). The results, displayed in Supplementary Figures 1, 2 and 3, illustrate that reproducibility is influenced by the spatial normalization pipeline regardless of the order in which the control task was carried out. Therefore, it is unlikely that the results obtained from our study reflect carry over effects from previous learning sessions.

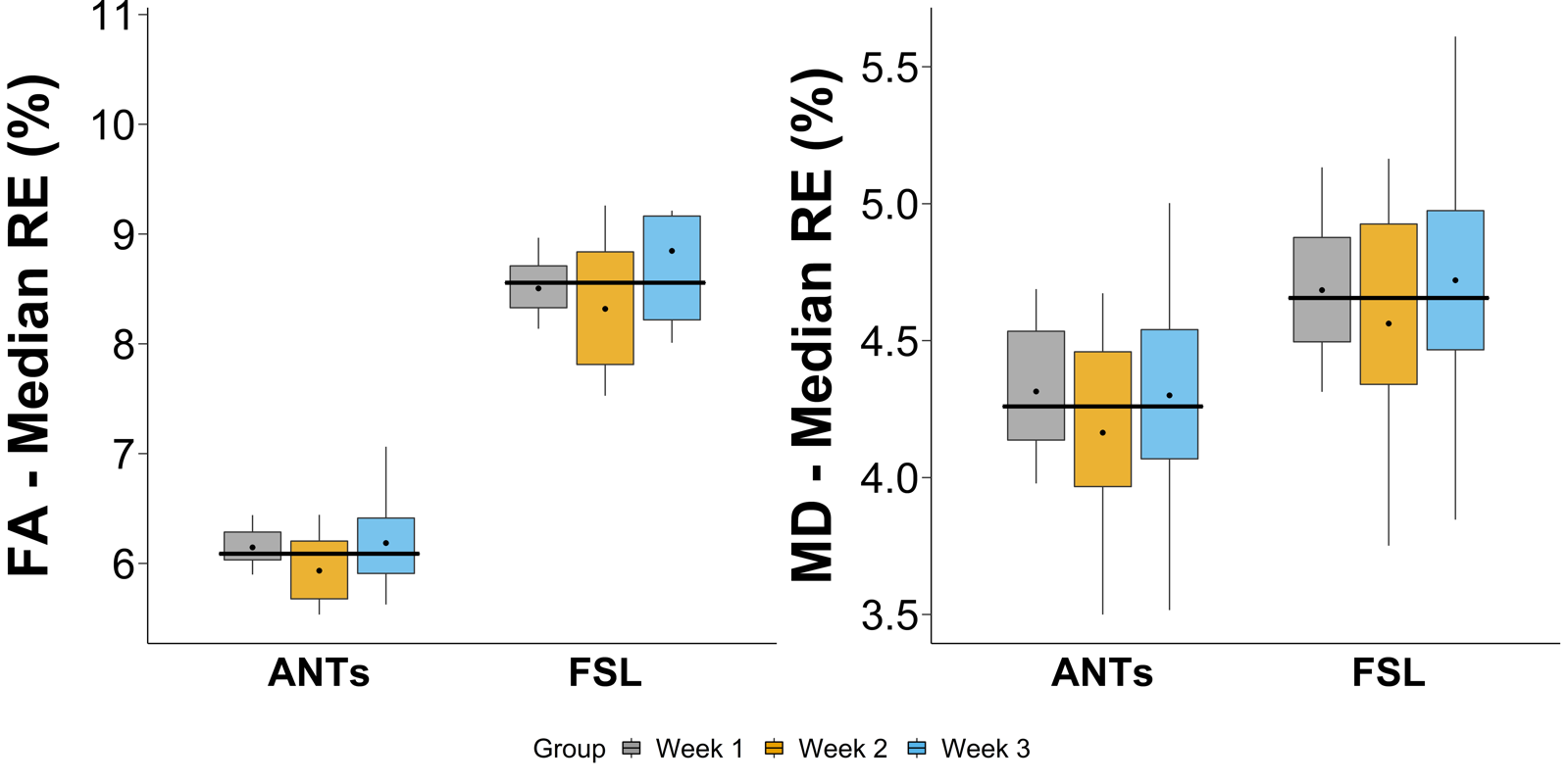

**Supplementary Figure 1:** Brain diffusion metric reproducibility effects of “Week” group and *Registration algorithm* (ANTs of FSL). Shown are the median RE obtained in white matter using ANTs or FSL for FA (left panel) and MD (right panel) for each Week. The dot illustrates the group mean. Wide black horizontal lines correspond to the mean of each pipeline across groups. The set of images used to calculate the reproducibility error (Week 1, Week 2 or Week 3) does not impact on results. Note that the Pipeline-by-Group interaction and main effect of Group terms of the two-way mixed ANOVA were not significant, neither for FA nor for MD (Pipeline-by-Group: F(2,17)=0.42, p=0.67 and Group: F(2,17)=0.46, p=0.64 for RE of FA; Pipeline-by-Group: F(2,17)=0.22, p=0.80 and Group: F(2,17)=0.26, p=0.78 for RE of MD). In contrast, the main effect of registration pipeline (ANTs vs FSL) was significant for both DTI measures (F(1,17)=1808.30, p<0.0001 for RE of FA; F(1,17)=449.78, p<0.0001 for RE of MD).

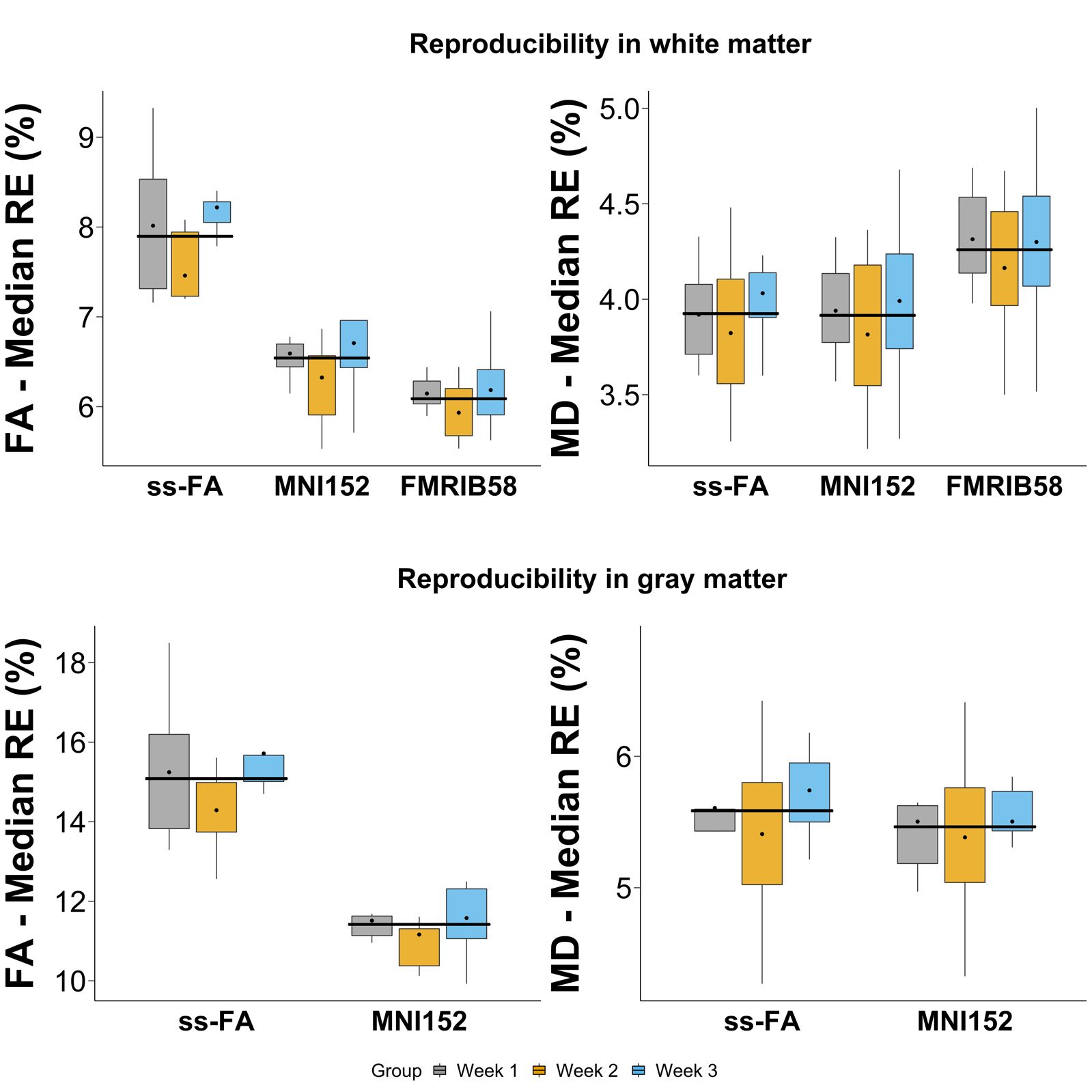
**Supplementary Figure 2:** Brain diffusion metric reproducibility effects of “Week” group and *Registration target* pipelines (MNI152 T1 template, FMRIB58 FA template or a study-specific FA template (ss-FA)). Shown are the median RE obtained in white matter (top) or gray matter (bottom) using MNI152, FMRIB58 or ss-FA for FA (left panel) and MD (right panel) for each Week. The dot illustrates the group mean. Wide black horizontal lines correspond to the mean of each pipeline across groups. The set of images used to calculate the reproducibility error (Week 1, Week 2 or Week 3) does not impact on results. The Pipeline-by-Group interaction and main effect of Group terms of the two-way mixed ANOVA were not significant, neither for FA nor for MD in WM (Pipeline-by-Group: F(2.53,22.79)=0.79, p=0.50 and Group: F(2,18)=1.98, p=0.17 for RE of FA; Pipeline-by-Group: F(2.63,23.64)=1.13, p=0.35 and Group: F(2,18)=0.40, p=0.68 for RE of MD). Yet, the main effect of Pipeline was significant for both DTI measures in WM (F(1.26,22.79)=114.01, p<0.0001 for RE of FA; F(1.31,23.64)=129.64, p<0.0001 for RE of MD). The Pipeline-by-Group interaction and main effect of Group terms of the two-way mixed ANOVA were not significant, neither for FA nor for MD in GM (Pipeline-by-Group: F(2,18)=0.94, p=0.41 and Group: F(2,18)=1.44, p=0.26 for RE of FA; Pipeline-by-Group: F(2,18)=0.50, p=0.61 and Group: F(2,18)=0.35, p=0.71 for RE of MD). The main effect of Pipeline was significant for FA in GM (F(1,18)=145.86, p<0.0001). On the other hand, the difference between pipelines was not significant for MD in GM F(1,18)=1.95, p=0.18).

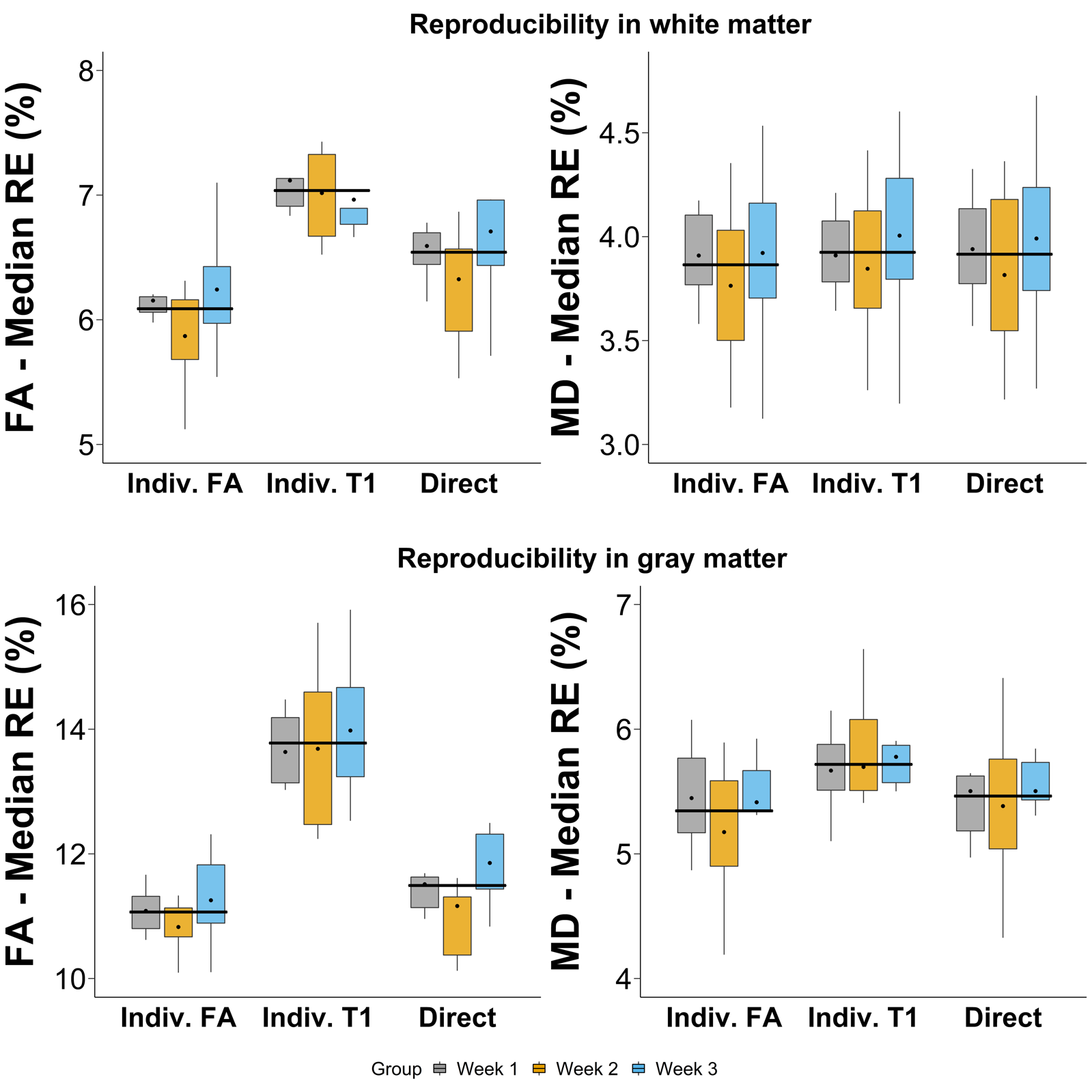
**Supplementary Figure 3:** Brain diffusion metric reproducibility effects of “Week” group and *Intermediate targets* pipeline used to warp DTI to MNI152 template. **Direct**: direct normalization, **Indiv. FA**: normalization via an individual FA template (created out of FA images from 9 scans per subject) and **Indiv. T1**: normalization via an individual T1 template (created out of T1 images from 9 scans per subject). Shown are the median RE obtained in white matter (superior row) and gray matter (inferior row) for FA (left panel) and MD (right panel) for each Week. The dot illustrates the group mean. Wide black horizontal lines correspond to the mean of each pipeline across groups. The set of images used to calculate the reproducibility error (Week 1, Week 2 or Week 3) does not impact on results. The Pipeline-by-Group interaction and main effect of Group terms of the two-way mixed ANOVA were not significant, neither for FA nor for MD in WM or GM

(Pipeline-by-Group F(4,32)=0.33, p=0.54, Group F(2,16)=0.99, p=0.39 for FA in WM; Pipeline-by-Group F(4,32)=0.64, p=0.64, Group F(2,16)=0.37, p=0.70 for MD in WM; Pipeline-by-Group F(4,32)=0.25, p=0.91, Group F(2,16)=0.51, p=0.61 for FA in GM; Pipeline-by-Group F(4,32)=0.65, p=0.63, Group F(2,16)=0.14, p=0.87 for MD in GM). Yet, the main effect of Pipeline was significant for both DTI measures in WM (F(2,32)=67.17, p<0.0001 for RE of FA; F(2,32)=6.91, p=0.003 for RE of MD) and in GM (F(2,32)=92.76, p<0.0001 for RE of FA; F(2,32)=21.39, p<0.0001 for RE of MD).
